# Transcriptional landscapes underlying Notch-induced lineage conversion and plasticity of mammary basal cells

**DOI:** 10.1101/2024.07.04.602034

**Authors:** Candice Merle, Calvin Rodrigues, Atefeh Pourkhalili Langeroudi, Robin Journot, Fabian Rost, Yiteng Dang, Steffen Rulands, Silvia Fre

## Abstract

The mammary epithelium derives from multipotent mammary stem cells (MaSCs) that progressively restrict their potency and engage into lineage commitment during embryonic development. Although postnatal mammary progenitors are lineage-restricted and unipotent, several lines of evidence have documented their extensive plasticity and ability to reactivate multipotency in several non-physiological contexts. We have previously shown that ectopic Notch1 activation in committed mammary basal cells, which never experience Notch activity in homeostatic conditions, triggers a progressive cell fate switch from basal to luminal cell identity in both the pubertal and adult mouse mammary gland. Here, we tested the conservation of this mechanism in other glandular epithelia and found that constitutive Notch1 signaling also induces a basal-to-luminal cell fate switch in adult cells of the lacrimal gland, the salivary gland, and the prostate. Since cells do not undergo lineage transition synchronously and this switch is progressive in time, we performed single cell transcriptomic analysis by SMART-Seq on index-sorted mutant mammary cells at different stages of lineage conversion, to reveal the molecular pathways underlying the fate transition. Combining single cell transcriptomics analyses with assays in organoid cultures, we demonstrate that proliferation of basal mutant cells is indispensable to convert them into luminal progenitors. We thus reveal the molecular mechanisms and individual transcriptional landscapes controlling lineage conversion and cellular plasticity of unipotent committed mammary cells *in vivo* with spatial and temporal resolution. Given the strong implications of Notch signaling in cancer, these results also provide important insights into the mechanisms that drive cellular transformation.

## Introduction

The adult mammary gland is composed of two epithelial layers featuring two main cell types: basal cells (BCs) in contact with the basement membrane and luminal cells (LCs) facing the ductal lumen, which can be further subdivided in ERα^pos^/PR^pos^ cells (also called Luminal Mature or LM) and ERα^neg^/PR^neg^ cells (often referred to as Luminal Progenitors or LP). It is now well-established that this tissue is maintained by unipotent lineage-restricted progenitors throughout adult life in homeostatic conditions, but these self-renewing committed cells retain a high degree of plasticity, as they can revert to multipotency in several circumstances.

This was first observed in transplantation experiments, when adult unipotent mammary cells displayed multilineage differentiation capacity and could generate a functional mammary gland composed of both BCs and LCs (Rodilla et al., 2015; Shackleton et al., 2006; Stingl et al., 2006; Van Keymeulen et al., 2011). Other experimental procedures involving the dissociation of BCs and LCs, including when they are grown separately in 3D organoid conditions, have shown reactivation of multipotency of BCs, that can also be induced by different types of epithelial injury causing tissue regeneration and, importantly, by oncogene activation (Jamieson et al., 2017; Jardé et al., 2016; Koren et al., 2015; Van Keymeulen et al., 2015). Moreover, recent elegant *in vivo* experiments illustrated the capacity of BCs to generate LCs upon genotoxic stress (Seldin and Macara, 2020)or LCs genetic ablation (Centonze et al., 2020), strongly suggesting that LCs restrict the default multipotency of BCs. We have previously shown that the binary fate choice between basal or luminal commitment is controlled by Notch signaling, a master regulator of cell fate choices in most vertebrate and invertebrate tissues, which is both necessary and sufficient for luminal fate specification in the mammary gland. Importantly, our previous studies uncovered that, besides its essential role in controlling fate decisions of embryonic multipotent mammary stem cells, constitutive and ectopic Notch activation in committed adult BCs, which never experience Notch activity, can also “reprogram” their lineage potential and induce their conversion into ERα^neg^/PR^neg^ luminal cells (Lilja et al., 2018).

The “reprogramming” capacity of mammary progenitors has important implications for cell differentiation as well as transformation; given that the cell fate switch did not occur in all cells at the same time, we set out to assess if the targeted BCs themselves transdifferentiate into LCs or if they respond to Notch activation by giving rise to luminal daughter cells. To this aim, we investigated the dynamics of the progressive lineage transition from basal to luminal fate to understand how the Notch-imposed cell fate switch is mechanistically achieved and to reveal the changes in transcriptional state of single Notch mutant cells *in vivo*, using two different Cre promoters and single cell RNA sequencing of index-sorted mutant cells at different stages of lineage transition. We found that the transcriptional changes associated with the transition from basal to luminal fate are progressive in time, triggered by the initial decrease in basal markers expression followed by the steady upregulation of luminal genes. Thanks to organoid cultures, we could also establish that proliferation is essential for cell fate conversion to occur, ruling out the possibility of a transdifferentiation event.

## Results

### Ectopic Notch activation induces a cell fate switch in four different bi-layered epithelia

We have previously shown that constitutive activation of Notch signaling through the ectopic expression of the ligand-independent, intracellular portion of the mouse Notch1 receptor (in R26-N1ICD-ires-nGFP gain-of-function mice) (Murtaugh et al., 2003) in mammary BCs, targeted by two different BC-specific inducible Cre promoters, SMACre^ERT2^ and K5Cre^ERT2^, is sufficient to trigger a progressive switch in cell fate and eventually forces all mutant cells to acquire a luminal Hormone Receptor^neg^ (HR^neg^) identity (Lilja et al., 2018).

Importantly, the *in vivo* “reprogramming” of the initially targeted BCs to LCs happens progressively over a long period of time (6 weeks), and nuclear GFP+ cells (nGFP^pos^), reporting Notch pathway activation, appear to transition through a phase resembling embryonic non-committed MaSCs, as revealed by their co-expression of basal (K14) and luminal (K8) markers (**Figure 1A-B**), prior to giving rise exclusively to fully committed luminal cells (**Figure 1A-B and Figure S1A**). This cell fate transition happens in unipotent adult BCs, as demonstrated by the exclusive labeling of α-SMA^pos^ BCs in control SMACre^ERT2^/mTmG mice, tracing the fate of targeted BCs and their progeny (**Figure S1B**).

**Figure 1.**
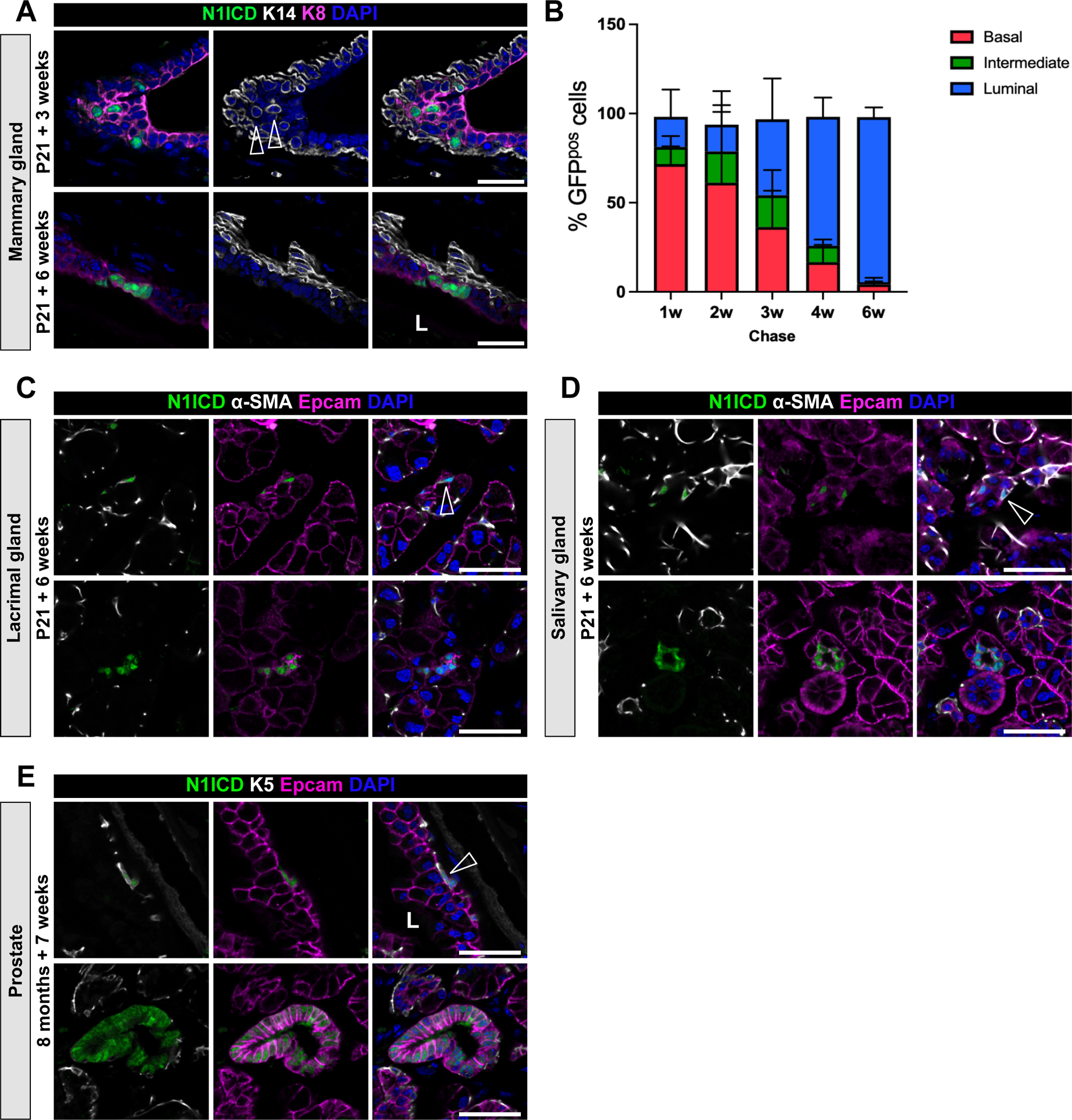
*In vivo* reprogramming of adult BCs to LCs by Notch1 activation in four bi-layered glandular epithelia. **A.** Representative sections of SMACre^ERT2^/N1ICD-ires-nGFP mammary glands induced at P21 and analyzed 3 or 6 weeks later by immunofluorescence for the basal marker K14 (white), the luminal marker K8 (purple) and nGFP (correlated to N1ICD expression in green). Nuclei are stained with DAPI. Empty arrowheads indicate mutant cells co-expressing nGFP, K14 and K8. **B.** Quantification by flow cytometry of the percentage of nGFP^pos^ basal (CD49f^high^/EPCAM^low^), intermediate (CD49f^med^/EPCAM^med^) and luminal (CD49f^low^/EPCAM^high^) cells 1, 2, 3, 4 and 6 weeks after tamoxifen induction at P21 (mean, SD, n). **C-D.** Representative sections of SMACre^ERT2^/N1ICD lacrimal glands (C) and salivary glands (D) induced at P21 and analyzed 6 weeks later by immunofluorescence for the basal marker α-SMA (white), the luminal marker Epcam (purple) and nGFP (N1ICD in green). Nuclei are stained with DAPI. **E.** Representative sections of K5Cre^ERT2^/N1ICD prostate induced at 8 months and analyzed 7 weeks later by immunostaining for the basal marker K5 (white), the luminal marker Epcam (purple) and nGFP (N1ICD in green). Scale bar represents 25µm in A, C-E. Empty arrowheads in C-E indicate cells that have not undergone cell fate switch at the time of the analysis. “L” indicates the lumen position in A and E.

Throughout the observed cell fate transition, hybrid cells co-expressing luminal and basal markers can be scored by flow cytometry analysis as dispersed cells laying between the luminal (EPCAM^high^/Cd49fl^ow^) and the basal (EPCAM^low^/Cd49^high^) gates (**Figure S1C, D**). The proportion of cells transiting through this intermediate state is highest mid-way through the lineage transition, around 3-4 weeks after Notch activation. Finally, the intermediate population completely disappears after a 6-week chase, when all mutant nGFP^pos^ cells have become luminal (**Figure 1B**, **Figure S1D**).

The Notch pathway is crucial for stem cell fate decisions in a variety of tissues. To assess the conservation of the role of Notch on binary cell fate choices in other glandular epithelia, we analyzed the plasticity and differentiation state of unipotent adult BCs induced to express N1ICD by the SMACre promoter (SMACre^ERT2^/N1ICD-nGFP) in the salivary gland and the lacrimal gland, and by the K5Cre promoter (K5Cre^ERT2^/N1ICD-nGFP) in the prostate. Remarkably, 6 weeks after N1ICD-nGFP ectopic expression, we found that most nGFP^pos^ cells had acquired a luminal identity also in the salivary and lacrimal gland, as well as in the prostate (**Figure 1C-E**), whereas control mGFP^pos^ BCs maintained their unipotency in SMACre^ERT2^/mTmG mice or K5Cre^ERT2^/mTmG mice in all examined tissues (**Figure S1E, F**). It is noteworthy that, although the vast majority of the targeted cells indeed switched to a luminal identity, we found a few mutant nGFP^pos^ salivary, lacrimal or prostate cells that maintained a basal phenotype after a 6-week chase, as assessed by their expression of α-SMA or K5. However, we decided not to further investigate the properties of these rare cells and we cannot therefore ascertain if they would also eventually turn into LCs after a longer time.

The striking conservation of the phenotype induced by ectopic Notch activation in four adult epithelia derived from different germ layers highlights the essential role of Notch signaling as a broad determinant of luminal cell fate.

### Intermediate mammary cells feature a hybrid transcriptional signature

The conspicuous robustness of the results we obtained using the same gain-of-function mutant mice in four different adult tissues demonstrates the high degree of plasticity of lineage-committed unipotent basal progenitors, that can readily change fate if homeostasis is perturbed, as previously observed in organoids (Jamieson et al., 2017; Jardé et al., 2016), in transplantation assays (Van Keymeulen et al., 2011) or, more recently, upon genotoxic agents exposure (Seldin and Macara, 2020) and in genetic ablation experiments *in vivo* (Centonze et al., 2020).

To gain mechanistic insights into the observed Notch-imposed cell fate switch and reveal the molecular pathways involved, we set out to decipher the properties of individual intermediate mammary cells at single cell level, to capture discrete gene expression states that could represent distinct differentiation trajectories. To this aim, we performed single cell RNA sequencing (scRNAseq) by SMART-SeqV2 on index-sorted mammary BCs, intermediate cells and LCs, both GFP^neg^ and GFP^pos^, isolated from pubertal mammary glands of SMACre^ERT2^/N1ICD and K5Cre^ERT2^/N1ICD mice, at different timepoints after Notch1 activation (1, 3, 4 and 6-week chase) (**Table 1**). Conditional expression of N1ICD was triggered in SMA^pos^ or K5^pos^ BCs, to assess if these two basal-specific Cre drivers would target different BCs characterized by distinct degrees of differentiation or specialization (Prater et al., 2014). Comparison of BCs “reprogramming” with the two Cre lines indicated that any BC can lineage convert to HR^neg^ LCs upon Notch ectopic activation, regardless of their differentiation status or plasticity (**Figure S2A)**.

**Table 1:** number of cells index-sorted per time and 96-well plate.

|  | Chase | LC GFP+ | BC GFP+ | Inter GFP+ | LC GFP- | BC GFP- |
| --- | --- | --- | --- | --- | --- | --- |
| SMACre/N1ICD | 1w | 6 | 73 | 11 | 6 |  |
| SMACre/N1ICD | 3w | 34 | 12 | 44 | 6 |  |
| SMACre/N1ICD | 3w | 30 | 34 | 26 | 6 |  |
| SMACre/N1ICD | 3w | 0 | 0 | 60 | 6 | 6 |
| K5Cre/N1ICD | 3w | 6 | 6 | 66 | 6 | 6 |
| K5Cre/N1ICD | 3w | 6 | 6 | 66 | 6 | 6 |
| SMACre/N1ICD | 4w | 22 | 17 | 51 | 6 |  |
| SMACre/N1ICD | 6w | 84 |  |  | 12 |  |

After data pre-processing, including quality control to remove cells of low quality, a total of 474 cells were subject to further analyses. Unsupervised clustering identified 5 distinct cell clusters that were composed of both mutant nGFP^pos^ and WT (nGFP^neg^) cells. One cluster was enriched for BCs, as confirmed by their high expression of *Krt5* and *Krt14*, and we called it BAS cluster. Two clusters were enriched for luminal markers: one representing luminal HR^neg^ cells, expressing *Krt8* and *Krt19*, that we termed HR^neg^ cluster, and the second one, mainly composed of WT luminal mature cells, expressing *Esr1* and *Pgr* coding for the Estrogen Receptor-α and Progesterone Receptors, named HR^pos^ (for Hormone Receptor positive) cluster. Interestingly, we identified two distinct clusters, called INT1 and INT2, that represented the intermediate cells that appear upon Notch activation (**Figure 2A-B, Figure S2B**).

**Figure 2.**
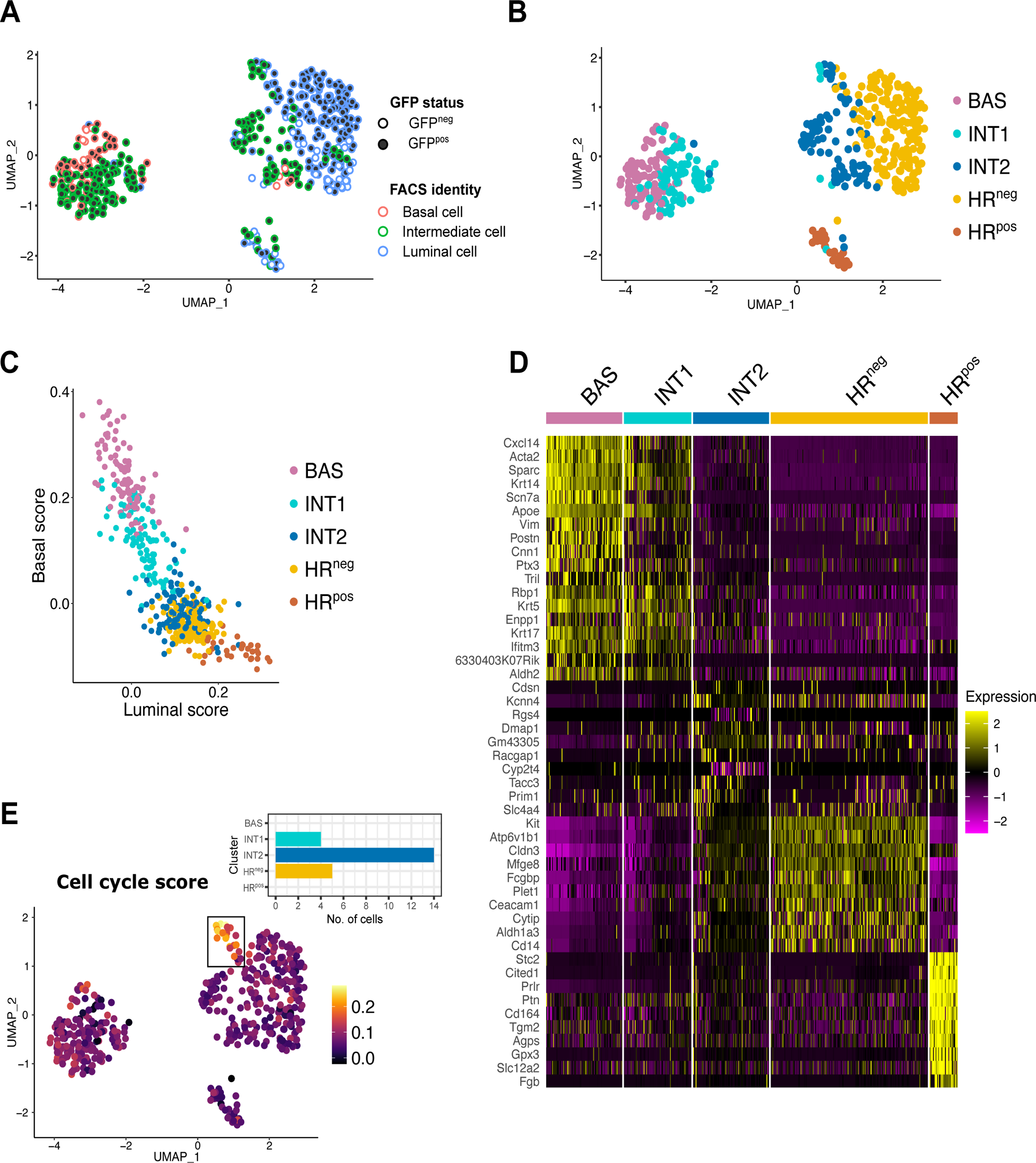
Index-sorted single cell RNAseq reveals the hybrid signatures of transitioning intermediate mutant cells. **A.** UMAP plot showing the identity of each index-sorted cell along with their GFP status. Cells are color-coded based on their FACS-defined identity as Basal (red), Intermediate (green) and Luminal (blue) cells. Mutant cells (GFP^pos^) are depicted as filled dots; WT cells (GFP^neg^) are shown as empty dots. **B.** UMAP plot showing clustering of single sequenced cells by Smart-seq2. 5 Seurat clusters were identified: BAS=Basal cells (pink), INT1=Intermediate 1 (turquoise), INT2=Intermediate 2 (dark blue), HR^neg^= Hormone Receptor^neg^ (yellow), and HR^pos^= Hormone Receptor^pos^ cells (brown). **C.** Plot representing the basal and luminal scores for each individual cell. Each dot represents a cell and their color corresponds to the clusters illustrated in (B). **D.** Heatmap of marker genes specific for each cell cluster illustrated in (B). Each column is color-coded according to the corresponding cell clusters from (B). The color key corresponds to normalized and scaled values of gene expression. **E.** UMAP plot showing enrichment for the GO term “Cell cycle score” across individual cells. The bar plot represents the number of cells, grouped by cluster, within the rectangular selected region in the UMAP.

We then calculated a basal and luminal score based on published transcriptomic profiles of adult Mammary Epithelial Cells (MECs) (Kendrick et al., 2008) (**Figure 2C, Figure S2C)**. As expected, both INT1 and INT2 clusters presented mixed basal and luminal scores, suggesting a hybrid signature characterized by the co-expression of luminal and basal markers.

Unsupervised cluster analysis of differentially expressed genes (DEGs) for each cluster (**Figure 2D**) revealed that the basal markers *Acta2*, *Sparc* and *Krt14* were strongly expressed in the BAS cluster and were progressively reduced in INT1, whereas genes typically associated with luminal identity, such as *Plet1*, *Kit* and *Aldh1a3*, were enriched in the HR^neg^ cluster and reduced in INT2.

However, we could not identify genes exclusive of the INT1 cluster and only 7 genes were specific of INT2. Moreover, these genes appeared to be upregulated only in few cells and did not define most of the cells in this cluster. Among these genes, we found *Tacc3*, involved in the stabilization of the mitotic spindle (Ding et al., 2017; Singh et al., 2014) and *Racgap1*, required for cytokinesis (Lekomtsev et al., 2012). Importantly, we also identified a group of cells, mainly belonging to the INT2 cluster, with a highly enriched cell cycle score (**Figure 2E**).

We also noticed that, although nGFP^pos^ and nGFP^neg^ LCs belonged to the same cluster, they appeared segregated based on their GFP status, suggesting that, even if they were scored as LCs by FACS, nGFP^pos^ mutant cells remained somehow different from fully differentiated LCs (**Figure 2A**). Consistent with this, Principal Component Analysis (PCA) confirmed that mutant LPs (GFP^pos^) at 3 and 4 weeks of chase do not entirely overlap with WT LPs (GFP^neg^) (**Figure S3A**). To reveal the differences between GFP^neg^ and GFP^pos^ LPs, we performed UMAP analysis exclusively within the LP cell cluster, and we could recognize two new clusters. Most GFP^neg^ cells are associated to one cluster, that we named luminal cells, and the other cluster was mainly composed of GFP^pos^ cells, that we called pre-luminal cells (**Figure S3B**). These two clusters are mainly distinguished by the time after N1ICD activation, as cells seem to acquire a pre-luminal identity at 1, 3 and 4 weeks and a more complete luminal identity after 6 weeks (**Figure S3C**). Although well-established luminal markers, such as *Krt18* and *Epcam*, were similarly expressed by these two clusters, other luminal genes, like *Trf* and *Clic6*, presented very low levels of expression in the pre-luminal cluster (**Figure S3D**), corroborating the notion that these cells, identified as luminal cells by cell sorting (based on EPCAM^high^ expression), have not yet entirely acquired a luminal identity.

Based on our computed basal and luminal scores and on the list of DEGs, the two clusters of intermediate cells expressed a mixed gene set between luminal and basal genes, with INT1 more closely related to the BAS cluster and INT2 more luminal, suggesting a progressive transcriptional switch from a basal to a luminal differentiation program.

Given that several single cell transcriptomic studies described a population of cells co-expressing basal and luminal markers (often called hybrid cells) in embryonic mammary glands (Giraddi et al., 2018; Pal et al., 2021; Wuidart et al., 2018) or in response to LCs ablation (Centonze et al., 2020), we wondered if the INT clusters we identified in our study reflected the presence of cells that reactivated embryonic or regenerative programs typical of multipotent MaSCs. To interrogate this, we compared our intermediate cells with hybrid cells identified in published datasets using Label Transfer from the Seurat package (Stuart et al., 2019), a variant of integration which allows the transfer of cluster labels from a reference dataset to a query dataset. The scRNAseq profile of mammary cells published by Wuidart and colleagues (Wuidart et al., 2018) comprised adult BCs and LCs, as well as “Embryonic Multipotent Progenitors” (EMPs) (CD49f^high^/Lgr5-GFP^high^) isolated from mammary tissue at embryonic day 14 (E14). EMPs co-expressed genes typical of BCs and LCs, but they also presented specific genes that were defined as the EMPs signature. By integrating our dataset with the Wuidart *et al*. dataset, we found, as expected, that the basal cell cluster (BAS) as well as the HR^neg^ and HR^pos^ clusters overlapped with the same adult cell types identified in that study (**Figure S4A**). Of interest, the INT1 and EMPs clusters co-localized in the PCA plots, validating their hybrid signature, whereas INT2 appears as a separate cluster that was not identified by Wuidart and colleagues. Using label transfer with Wuidart *et al*. as a reference and visualizing the transferred labels with an alluvium plot, we found that the INT1 cluster is associated with both the BAS and HR^neg^ clusters whereas INT2 is almost entirely linked to the HR^neg^ cluster and does not resemble to the BAS cluster (**Figure S4B**). To our surprise, however, this comparative analysis indicated that the EMPs signature was not specifically expressed in INT1 or INT2 clusters (**Figure S4C**). This comparative analysis indicated that the intermediate cells that we identified in our dataset do not necessarily revert to an embryonic multipotent progenitor state similar to the EMPs sequenced at embryonic day E14 by Wuidart and colleagues.

We then integrated our dataset with the sequencing results from Centonze *et al*. (Centonze et al., 2020), who performed scRNAseq of adult BCs and LCs following genetic ablation of a fraction of LCs *in vivo*. This genetic intervention induced reactivation of multipotency in BCs and the appearance of a population of hybrid cells (referred to as “Hybrid”) showing co-expression of basal and luminal markers. PCA of these integrated datasets indicated that our INT1 cluster closely integrates with the Hybrid cells and partially overlaps with the adult BAS cluster, whereas, once again, the INT2 cluster represent a separate cluster that shares lower resemblance with hybrids cells sequenced by Centonze and colleagues (**Figure S4D**). The analysis of the same dataset using the alluvium plot representation confirmed that our INT1 cluster is more transcriptionally similar to BAS and Hybrid cells from *Centonze et al.*, whereas INT2 cells appear more closely related to HR^neg^ cells (**Figure S4E)**.

This comparative *in silico* analysis suggests that the INT1 cluster represents a hybrid transcriptional cell state in between basal and luminal lineages, similarly to the Hybrid cluster reported by Centonze *et al*. On the contrary, the INT2 cluster is uniquely found upon Notch activation and, as such, it may represent a distinctive cluster, possibly more related to committed luminal cells.

In conclusion, the integration of our single cell profiles with published datasets indicates that the intermediate cells that appear upon Notch ectopic activation, although featuring a mixed signature, are different from embryonic multipotent MaSCs, and consequently they represent an adult hybrid state, likely denoting a transiting stage between BCs and LCs.

### Transcriptional landscapes underlying the progressive lineage transition from BCs to LCs

To examine the gradual transcriptional changes that occur during the cell fate switch from BCs to LCs, we then performed a slingshot trajectory analysis within the PCA space, denoting the BAS cluster as the origin. Interestingly, we observed a forked pattern presenting 2 separate trajectories (**Figure 3A**). Both trajectories passed through the INT1 and INT2 clusters, but one path terminates in the HR^neg^ cluster (Trajectory 1) while the other ends with the HR^pos^ cluster (Trajectory 2). The divergence of the two trajectories was observed around the INT2 cluster, suggesting that the two luminal identities are specified at the latest intermediate stage.

**Figure 3.**
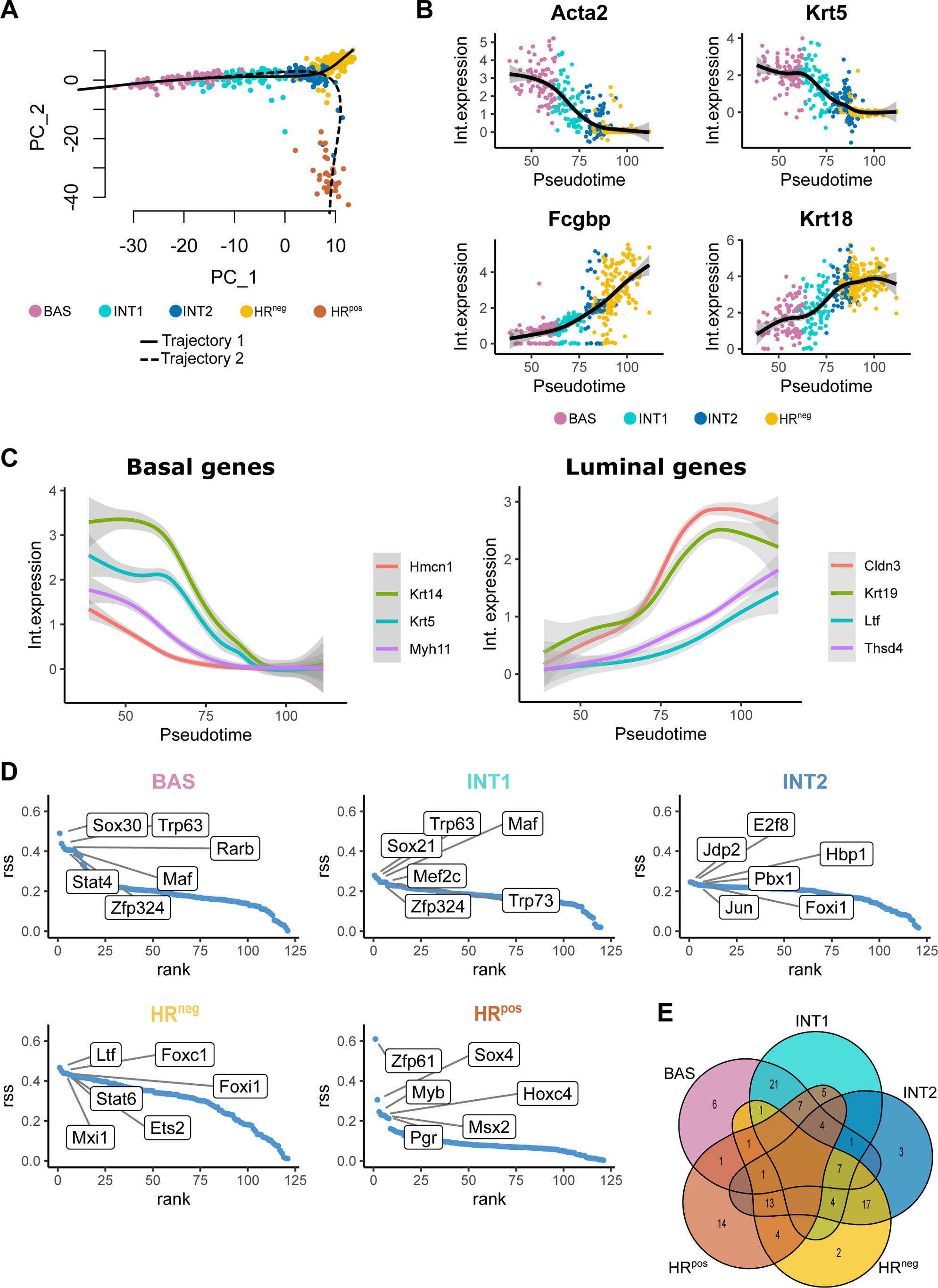
Cell trajectory and transcriptional signatures defining the progressive transition from basal to luminal identity. **A.** Slingshot trajectory analysis showing two cellular paths, connecting BAS cells to HR^neg^ (trajectory 1) or HR^pos^ (trajectory 2) clusters in a PCA plot. **B.** Expression of selected genes within cells plotted along trajectory 1 in pseudotime. The integrated gene expression is plotted; dots correspond to individual cells color-coded according to the UMAP clusters from Fig. 2B. **C.** Expression of selected basal and luminal genes along pseudotime trajectory 1. **D.** SCENIC analysis showing the Regulon specificity score (RSS) for each cluster: only the 6 most significant TF regulons showing cluster-specific activity are indicated. **E.** Venn diagram presenting the number of overlapping regulons among the 50 most significant TF regulons for each cell cluster.

Given that Notch1 activity is restricted to ERα^neg^/PR^neg^ luminal cells and that the cell fate switch induced by Notch activation eventually converts the targeted BCs exclusively into ERα^neg^/PR^neg^ LCs, we then focused our analysis on trajectory 1 (BAS to HR^neg^). For this, we used Tradeseq, which performs Generalized Additive Models (GAM) fitting to the gene expression variation along a pseudotime, detecting genes which significantly vary along the pseudotime. During the transition from BAS to HR^neg^ clusters, we observed the previously detected trend of progressive decrease in expression of classical basal markers, such as *Acta2* and *Krt5,* and gradual increase in luminal gene expression, including luminal markers like *Krt18* and *Fcgbp* (**Figure 3B, Figure S5A**). In general, most genes followed a trend of constant increase or decrease of expression during the progressive switch from basal to luminal cell identity. Along pseudotime, we noticed that the first event detectable at the transcriptional level consists in the downregulation of basal genes, as we had previously found in bulk RNAseq experiments (Lilja et al., 2018), and this is associated with cells belonging to the INT1 cluster. Later on along the pseudotime, the expression of luminal markers kicks in, in cells belonging to the INT2 cluster, which present co-expression of several genes typical of either BCs or LCs, such as *Krt19* and *Cldn3*, most likely corresponding to the K14/K8 double positive cells that we observed by immunostaining (**Figure 1A**) (Lilja et al., 2018). Finally, the last step of the transition is characterized by the complete loss of basal genes and the steady increase of expression of luminal lineage genes (**Figure S5A**).

The pseudotime analysis we performed clearly indicates the progressive nature of the lineage transition induced by ectopic Notch1 activation, characterized by continuous and gradual transcriptional changes underlying the sequential change in cell identity. We thus conclude that lineage conversion requires a stepwise and asynchronous change in transcriptional programs, with some basal genes downregulated early and others that take a longer time to be repressed. For example, the gene *Hmcn1*, encoding the immunoglobin superfamily member Hemicentin 1 and the smooth muscle myosin heavy chain *Myh11*, a well-described marker of the basal lineage (Prater et al., 2014), are among the earliest basal genes to be downregulated, whereas the typical basal cytokeratins *Krt14* and *Krt5* decrease in expression later (**Figure 3C**). The same is true for acquisition of a luminal identity, with genes such as *Krt19* and *Cldn3* that are upregulated very early during the transition, and others, like *Ltf* and *Thsd4*, whose upregulation is only observed toward the end of the transition in pseudotime (**Figure 3C**).

This temporal analysis along the pseudotime indicates that the transition from basal to luminal identity is a long and progressive process, involving the initial repression of basal genes, and subsequently the gradual activation of expression of luminal genes.

### Gene regulatory network analysis uncovers the molecular signatures of transitioning cells

To further capture the regulatory mechanisms at work during the cell fate switch, and to identify potential transcriptional nodes that could represent general regulators of cell plasticity, we then performed SCENIC analysis on our dataset. The SCENIC algorithm examines the activity of transcription factor regulons, consisting of transcription factors and their targets, within individual cells (Aibar et al., 2017). We used the Regulon Specificity Score (RSS) to identify regulons showing enriched activity in each cell cluster. We observed, as predicted, elevated activity of the Progesterone Receptor (*Pgr*) regulon in HR^pos^ cells and of *Foxc1* in HR^neg^ luminal progenitors, consistent with a previous report (Sizemore et al., 2013) (**Figure 3D**). Among the top 50 regulons enriched in cluster INT1, we could not pinpoint any that was exclusive for this cluster and was not shared with other clusters, and most of these regulons were found in both INT1 and BAS clusters, indicating a strong similarity of INT1 cells with basal cell identity (**Figure 3E**). Likewise, most of the regulons enriched in cluster INT2 were shared with the HR^neg^ cluster. This analysis corroborates our findings indicating that the early steps of lineage switch involve suppression of basal regulons, such as *Trp63* and *Trp73*, followed by the steady and progressive increase of activity of luminal-specific regulons, such as *Jun* or *Stat6* (**Figure S5B**). Consistent with our results, when we analyzed DEGs corresponding to each cluster, we could not identify regulons that would be unique INT1 cells. We could however find 3 regulons, *Brca1*, *E2f8* and *E2f1*, which were specific to INT2 cells, and these are all linked to elevated proliferation. Interestingly, these regulons are specifically enriched in highly proliferative cells belonging to the INT1, INT2 and HR^neg^ clusters (**Figure S5C**).

This analysis suggests that the lineage conversion involves activation of a proliferative signature, particularly relevant in cells belonging to the INT2 cluster, for mutant cells to complete the fate transition and engage into the transcriptional program characteristic of luminal cells.

### Proliferation is indispensable for switching cell identity

The scRNAseq analysis on individual mutant cells undergoing the cell fate transition allowed us to identify a group of cells, mainly belonging to the INT2 cluster, that presents a high cell cycle score and an upregulated activity of regulons linked with active proliferation. In addition, we have triggered Notch activation both at puberty and in adult mice and found that adult mammary cells take much longer to complete the transition from basal to luminal identity. While induction before puberty (at postnatal day P21) results in all mutant nGFP^pos^ cells to become LPs within 6 weeks, in adult mice, where cell divisions are less frequent, the complete switch is achieved in 10 weeks (data not shown). Based on these results, we formulated the hypothesis that the switch in cell identity does not simply represent a transdifferentiation event, bypassing cell division, but rather requires actively proliferating cells that respond to Notch activation by giving rise to luminal daughter cells.

In order to experimentally test if proliferation was required for the transition from BCs to LCs induced by ectopic Notch1 activation, we thus implemented the culture of 3D mammary organoids (Charifou et al., 2021; Jardé et al., 2016). First, we established that this *in vitro* system was suitable to study the cell fate switch, by demonstrating that WT cells derived from the adult mammary epithelium maintain their unipotent behavior in the organoid culture conditions (**Figure S6A**). Indeed, lineage tracing of WT BCs in organoids, using SMACre^ERT2^/mTmG mice (Muzumdar et al., 2007), revealed that exclusively basal daughter cells were derived from the initially labelled BCs and were therefore marked by our lineage tracer membrane mGFP. Importantly, upon *in vitro* Notch1 activation via 4-hydroxitamiofen (4-OHT) administration to the organoid medium, we could recapitulate the progressive transition of mutant nGFP^pos^ cells from the basal to the luminal lineage, correlated with increased expression of K8 and loss of α-SMA, robustly reflecting the data acquired *in vivo* (**Figure 4A**). Remarkably, a complete cell fate switch in organoids could be achieved within only 6 days after induction of N1ICD expression (**Figure S6B**). Thus, the organoid system allows us to induce a rapid cell fate switch, much faster than *in vivo*, and to target more cells, such that some organoids were exclusively composed of luminal nGFP^pos^ cells 6 days after Cre induction (**Figure 4A**).

**Figure 4.**
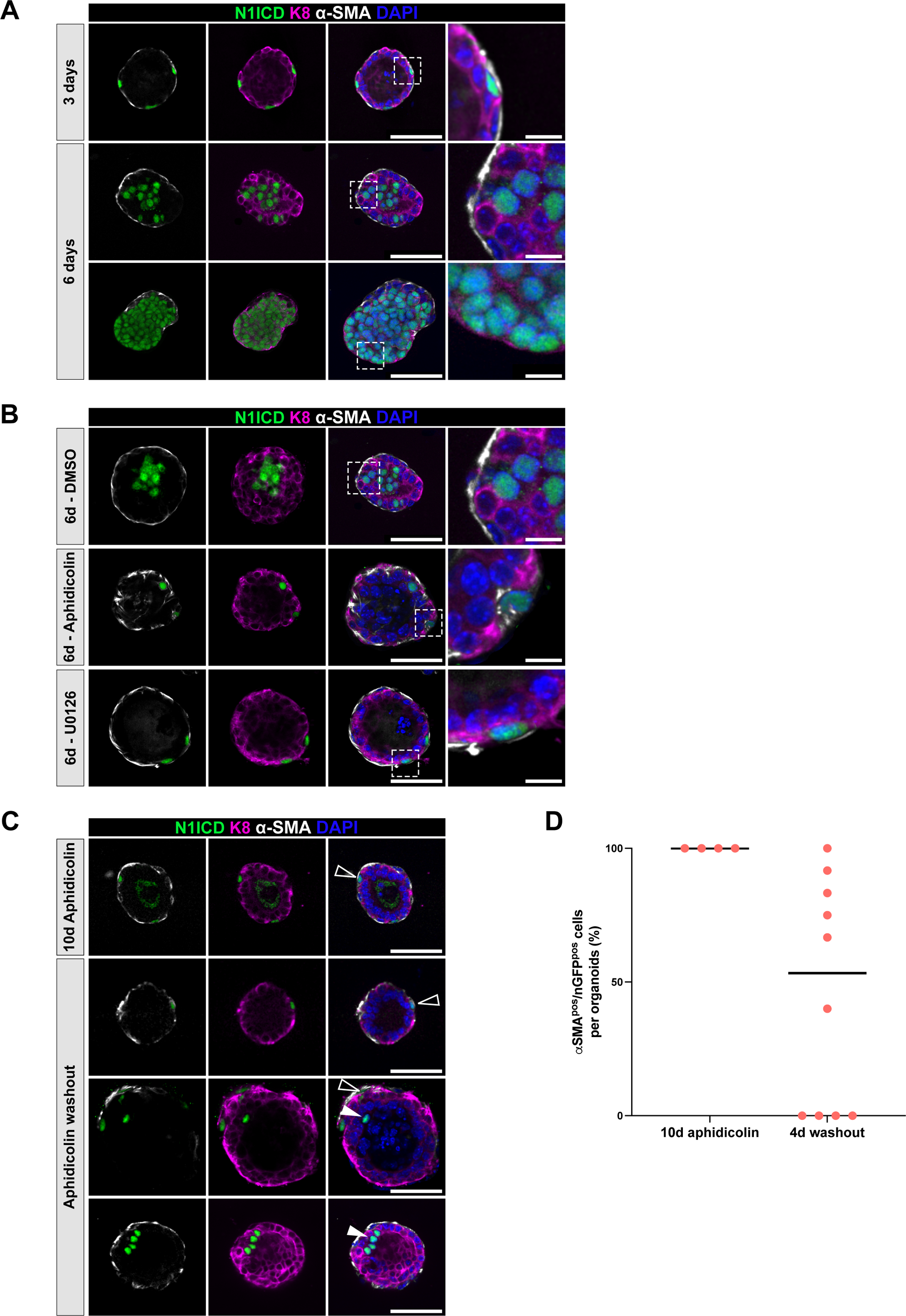
Proliferation is an obligatory step for lineage transition to occur in organoids. **A-C.** Representative images showing immunofluorescence for nGFP (N1ICD in green), luminal K8 (purple) and basal α-SMA (white) expression in SMACre^ERT2^/N1ICD mutant organoids 3 or 6 days after 4-OHT induction in (A); in SMACre^ERT2^/N1ICD mutant organoids treated with DMSO, Aphidicolin or U0126 in (B) and in SMACre^ERT2^/N1ICD mutant organoids treated with Aphidicolin for 6 days and grown for another 4 days upon Aphidicolin washout or treated with Aphidicolin for 10 consecutive days. Nuclei are stained with DAPI in blue. Scale bar represents 50µm in A-C and 10 µm (in A-B) for the magnified insets. Empty arrowheads indicate cells that have not undergone cell fate switch at the time of the analysis, white arrow heads indicate nGFP^pos^ luminal cells. **D.** Quantification of the proportion of basal nGFP^pos^ mutant cells within each organoid after 10 days of aphidicolin or after Aphidicolin washout for 4 days. The trait indicates the mean value.

Our *in vivo* data, corroborated by the single cell transcriptional analysis at different time points after Notch activation, demonstrated that the lineage transition is asynchronous, with some cells switching to a luminal fate more rapidly than others, thus indicating a heterogeneous competence of different targeted BCs to readily respond to Notch activation. Given the fact that resting adult mammary cells take longer to switch than proliferating pubertal cells and that instead organoids take less time, we then investigated the involvement of proliferation in dictating the differential readiness of BCs to transition towards a luminal fate. Validating our hypothesis, pharmacological block of proliferation in organoids, by supplementing the medium with either Aphidicolin, an inhibitor of DNA polymerase, or U0126, an inhibitor of MAPK activation, confirmed by Edu staining (**Figure S6C-D**), resulted in a complete arrest of the cell fate switch (**Figure 4B**), contrary to DMSO-treated control organoids, where the lineage transition of mutant nGFP^pos^ cells was completed within 6 days. To confirm that the observed block of cell fate switch was indeed directly associated with proliferation arrest, we then removed Aphidicolin after 6 days of treatment. Four days after washout, we found that 53% of the nGFP^pos^ cells re-entered the fate transition program and became luminal within 6 days (**Figure 4C, D**). It is noteworthy that the heterogeneous behavior of mutant nGFP^pos^ cells could be observed even within the same organoid, with some mutant cells readily switching to luminal fate upon aphidicolin washout and others more refractory to enter the lineage transition (**Figure 4C, D**). This experiment demonstrated that the arrest in cell fate switch can be reversed, and that proliferation is an obligatory step for lineage conversion.

We then assessed the temporal dynamics of the fate transition in organoids, and for this we used SMACre^ERT2^/mTmG/N1ICD compound mice, allowing us to track mutant cells following the fluorescence of membrane-tagged GFP (mGFP from the mTmG allele) in real time. In fact, the nGFP expressed with the N1ICD allele is not detectable by live microscopy, as it requires immunostaining with anti-GFP antibodies. In this experimental setting, we could indeed identify and track by time-lapse microscopy mGFP^pos^ cells, initially localized in the basal compartment, that enter cell cycle and subsequently move to a luminal internal position (**Figure 5A**). Some mGFP^pos^ cells instead remained in the basal compartment after mitosis, undoubtedly representing WT BCs that only floxed the mTmG reporter but not the more refractory N1ICD allele (**Figure 5B**). Given the observed lack of complete overlap between mGFP^pos^ cells (from the neutral mTmG allele) and N1ICD-expressing mutant cells, we performed immunostaining for the Notch1 direct target Hes1 and confirmed that all mGFP^pos^ cells that converted to a luminal identity were indeed mutant, whereas the ones that remained BCs did not present Notch activation, as assessed by Hes1 protein expression (**Figure 5C**).

**Figure 5.**
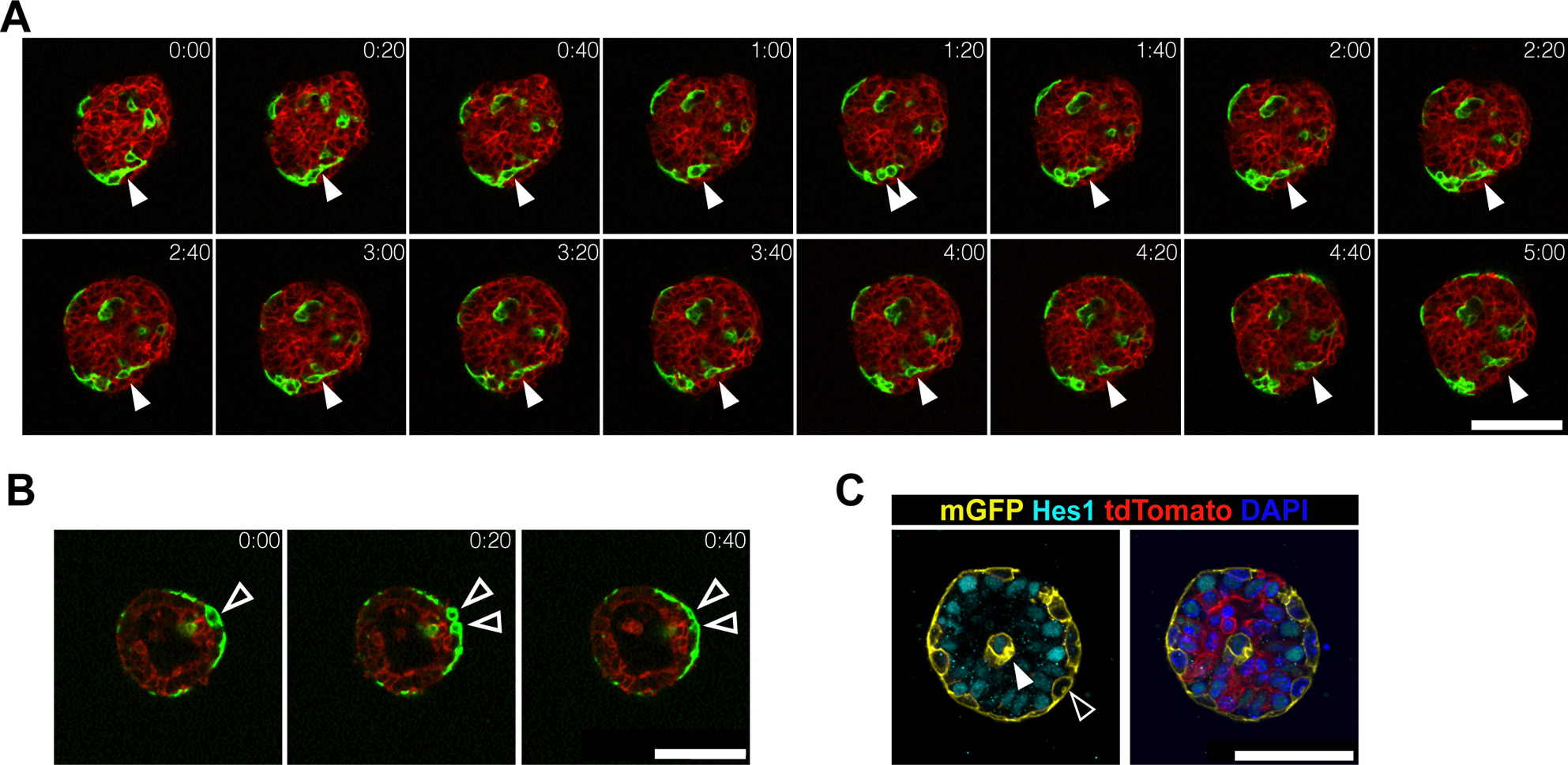
Dynamic behavior of lineage transitioning cells by time-lapse analysis. **A-B.** Sequential time-lapse images of SMACre^ERT2^/mTmG/N1ICD organoids showing recombined GFP^pos^ (green) cell rearrangements over 5 h. Red: non-recombined tdTomato^pos^ cells. White arrowheads in (A) pinpoint a mutant BC that first divides (between 1h 20min and 1h 40 min time frames) and then one of the two daughter cells that moves to a luminal position after mitosis. The empty arrowheads in (B) depict the mitosis of a WT basal cell whose daughters stay in the basal outer cell layer after division. Scale bar 25µm. **C.** Representative images showing immunofluorescence for mGFP (indicating recombined cells in yellow), Hes1 (marking nuclei and reflecting Notch activation in turquoise) and tdTomato expression in red in organoids grown for 3 days. Nuclei are stained with DAPI in blue. White arrowheads indicate mutant cells (Hes1 positive) and black arrowhead indicate WT cells (Hes1 negative). Scale bar 50µm.

These results demonstrate that the lineage switch induced by Notch1 is achieved through a progressive change in cell identity, whereby mutant cells transit through an intermediate metastable state, that requires their capacity to enter mitosis.

## Discussion

We report here that Notch signaling is a gatekeeper of luminal cell fate and that this critical role of dictating binary cell fate choices is conserved in several tissues, as demonstrated by the fact that ectopic Notch1 activation in committed adult BCs reprograms them toward a luminal identity in four different glandular epithelia. Importantly, we observed both *in vivo* and in organoids that BCs ectopically induced to activate Notch signaling rapidly move towards the ductal lumen, while acquiring luminal characteristics, indicating that intrinsic signals dictate cell fate, leading to positional changes and rearrangements of cells within bi-layered branched epithelia. Future studies will be required to probe if Notch activity directly influence cell position and movements or if other factors act on cell dynamics and contribute to establish the definitive commitment toward a luminal cell fate.

We demonstrate here that the transition from basal to luminal state is achieved through a progressive transcriptional switch, triggered by the initial downregulation of basal genes, followed by upregulation of luminal differentiation programs. These two cellular states are not exclusive, as demonstrated by the presence of hybrid cells co-expressing luminal and basal markers (K14^pos^/K8^pos^ cells). While the presence of hybrid cells has been reported in several contexts and it is believed to reflect the remarkable cellular plasticity of mammary BCs, it was not known that proliferation is a mandatory step to induce this intermediate metastable cell state and to accomplish the lineage switch to LCs. These findings indicate that adult mammary BCs, when forced to activate Notch signaling and change fate, do not undergo transdifferentiation, but rather that they are reprogrammed to a plastic state that, despite their initial unipotency, enables them to give rise to LCs, thus alters their differentiation potential independently of their position within the tissue. Mutant BCs appear to transition through an intermediate transient phase of co-expression of basal and luminal markers before attaining a luminal identity and eventually giving rise exclusively to fully ‘reprogrammed’ LCs. We show that BCs initially reduce the expression of basal genes, and then they enter a state of active proliferation, which results in the generation of luminal daughter cells. This behavior reflects their extensive plasticity and does not necessarily require Notch activity, since WT BCs induced to reactivate bipotency by LCs genetic ablation, also require proliferation to give rise to new luminal daughters, as shown by the fact that decreasing proliferation using a CDK1 inhibitor or by p21 overexpression in mammary organoids impaired BCs multipotency (Centonze et al., 2020). The hybrid cell state, characterized by co-expression of basal and luminal genes within the same cell, can also be found in breast cancer and it is often associated to a multipotency state (Van Keymeulen et al., 2015; Koren et al., 2015). However, the continuous expression of active Notch1 in our model prevents the maintenance of multipotent cells since cell differentiation is biased toward a luminal fate and eventually all mutant cells become HR^neg^ progenitors.

Differentiation and cell cycle are usually two cellular anti-correlated processes. However, the ectopic activation of Notch1 in differentiated BCs could induce the expression of cytokine-related cell cycle genes, like CDK1 (Ronchini and Capobianco, 2001), and at the same time activate transcription factors related to luminal differentiation, that we observed through the early activation of Jun or Foxi1, or recruit chromatin modifiers that could potentially tilt the balance of activation/repression on bivalent lineage promoters.

We report here that ectopic Notch activation results in the reprogramming of BCs into HR^neg^ progenitors. Given that *in vivo* Notch activation in BCs is mosaic, we do not observe an overt phenotype at the tissue level, since WT BCs that escaped tamoxifen induction can compensate for the mutant BCs that are lost to give rise to LCs. However, in organoids we could document the clonal expansion of mutant N1ICD-expressing LCs, that appear to gain a competitive advantage and eventually can form organoids composed exclusively of mutant nGFP^pos^ luminal cells (**Fig. 4A**, lower raw). Even if these mutant cells cluster close to WT LCs by UMAP analysis (**Figure 2A**), it is well established that Notch gain-of-function mice can form mammary tumours (Bouras et al., 2008; Callahan and Smith, 2000; Diévart et al., 1999). Indeed, deregulated Notch activation has been shown to induce mammary carcinomas (Diévart et al., 1999) and to affect human mammary cell transformation (Stylianou et al., 2006), stem cell maintenance (Harrison et al., 2010) and to be associated with poor outcome in breast cancer patients (Reedijk et al., 2005). Moreover, we found that constitutive Notch1 activation both when targeted to all mammary cells (with MMTV-Cre) and when restricted to HR^neg^ (with N1Cre^ERT2^ mice) results in pregnancy-dependent mammary hyperplasia (our unpublished observations). Of interest, our preliminary analyses suggest that the LPs generated by ectopic Notch1 activation in BCs (with both SMACre^ERT2^ and K5Cre^ERT2^) are also susceptible to transformation, and they promote the growth of hyperplastic lesions upon successive rounds of lactation and involution. These results carry important implications in breast cancer, revealing that Notch signaling is not only required for specifying luminal progenitor cells in the normal mammary gland, but that sustained and aberrant Notch activation in differentiated and lineage-committed cells has the potential to promote the appearance of mammary tumors, given its paramount role in the control of the delicate equilibrium between differentiation and proliferation that is necessary for healthy tissue homeostasis. These observations reinforce the concept that the mechanistic processes through which stem cells commit to a particular differentiation path mirror those hijacked by oncogenes to trigger cellular transformation across various tissues (Blanpain and Fuchs, 2014). Therefore, unraveling these mechanisms is crucial for better comprehending the genesis of cancer.

## Material and Methods

### Mice

SMA-Cre^ERT2^ (Wendling et al., 2009) and K5-Cre^ERT2^ (Indra et al., 1999) were crossed with a conditional gain-of-function Notch1 mutant mouse (Rosa-N1ICD-IRES-nGFP) (Murtaugh et al., 2003) or with the double fluorescent reporter Rosa26^mT/mG^ (Muzumdar et al., 2007). Reporter expression was induced by intraperitoneal injection of tamoxifen (1mg/10g of weight) at postnatal day P21.

### Ethics Statement

All studies and procedures involving animals were in agreement with the recommendations of the European Community (2010/63/UE) for the Protection of Vertebrate Animals used for Experimental and other Scientific Purposes. Approval was provided by the ethics committee of the French Ministry of Research (reference APAFIS #34364-202112151422480). We comply with internationally established principles of replacement, reduction, and refinement in accordance with the Guide for the Care and Use of Laboratory Animals (NRC 2011). Husbandry, supply of animals, as well as maintenance and care in the Animal Facility of Institut Curie (facility license #C75–05–18) before and during experiments fully satisfied the animal’s needs and welfare. All mice were housed and bred in a specific-pathogen-free (SPF) barrier facility with a 12:12 hr light-dark cycle and food and water available *ad libitum*. Mice were sacrificed by cervical dislocation.

### Mammary gland dissociation and cell sorting

Mammary glands were harvested and digested with collagenase (Roche, 57981821, 3mg/ml) and hyaluronidase (Sigma, H3884, 200U/ml) for 90min at 37°C under agitation. Following washes, cells were dissociated with Trypsin for 1min, dispase for 5min (200U/ml) and DNAseI (D4527, Sigma-Aldrich, 200U/ml) and then filtered through a 40µm cell strainer to obtain a single cell preparation. Cells were incubated for 30min with the following antibodies in 1:100 concentration: APC anti-mouse CD45 (Biolegend), APC anti-mouse Ter119 (Biolegend), APC anti-mouse CD31 (Biolegend), PE anti-mouse Epcam (Biolegend), APC-Cy7 anti-mouse CD49f (Biolegend). Single cell preparation was resuspended in flow buffer containing PBS, EDTA 5mM, BSA 1%, FBS 1% and DAPI. Dead cells (DAPI^pos^) and Lin^pos^ non-epithelial cells were excluded before analysis using FACS ARIA flow cytometer (BD). The results were analyzed using FlowJo software. Single sorted cells were deposited in 96-well plates containing SUPERase-In RNase Inhibitor (20U/µl, Sigma, AM2694), 10% Triton X-10 and DEPC-treated H_2_0 to library preparation using Smart-Seq2 protocol.

### Immunofluorescence on OCT sections

Mammary, salivary and lacrimal glands and prostates were harvested and fixed at room temperature in PFA 4% for 1h. Tissues were incubated for 3 days at 4°C in sucrose 30% and embedded in Optimal Cutting Temperature (OCT). Immunostainings were performed with 10µm sections. Antibodies used were rabbit anti-GFP (Institut Curie antibody platform, 1/300e), rat anti-K8 (TROMA-1, DSHB, 1/300e), chicken anti-K14 (906004, Biolegend, 1/500e), anti-αSMA coupled with AF488 (clone 1A4, F3777, Sigma-Aldrich) and chicken anti-K5 (905901, BioLegend). Fluorochrome-conjugated secondary antibodies included Cy5-conjugated anti-rat IgG (A102525, Invitrogen), Cy3-conjugated anti-rabbit IgG (A10520, Invitrogen) AlexaFluor488 anti-chicken IgG (A11039, Invitrogen) and AlexaFluor 488-conjugated anti-rabbit IgG (A21206, Invitrogen).

### Organoids culture

Primary mammary organoids were prepared from 2 to 3-months-old female mice. Mammary glands were collected, pooled and chopped to approximately 1-mm^3^ pieces and proceed to enzymatic digestion with 2 mg/mL collagenase A and 2 mg/mL trypsin for 30min at 37°C under agitation. Then, pieces were exposed to five rounds of differential centrifugation at 500g for 15 seconds in order to remove stromal cells. The organoids were resuspended in DMEM/F12 supplemented with 1× insulin–transferrin–selenium supplement, 100 U/mL of penicillin and 100μg/mL of streptomycin.

Organoids were resuspended in Matrigel® (Corning) and plated at 200 organoids for 30µl of Matrigel in 24-well plate. Matrigel drops were covered by culture medium and incubated at 37°C with 5% CO2. Activation of N1CD was triggered by 4-OHT (200nM) added to culture medium for 24h. To block proliferation U0126 (5 µM) or Aphidicolin (0.6µM) were added to culture medium for 6 to 10 days. Medium was changed every two days.

### Organoids staining

For immunostaining, organoids were fixed in 4% PFA for 10 min at room temperature, followed by 1h of permeabilization (1% Triton in PBS) and 2h incubation with blocking buffer (0.25% Triton / 2% BSA / 5% FBS / PBS). Primary antibodies were incubated overnight at 4°C and secondary antibodies and DAPI for 5h at room temperature. Antibodies used were anti-K8 (TROMA-1), rabbit anti-αSMA (NB600-531, Novus Biologicals), anti-α-SMA coupled to FITC (clone 1A4, F3777, Sigma-Aldrich), rabbit anti-GFP (Institut Curie antibody platform), rabbit anti-Hes1 (11988, Cell Signaling) and secondary antibodies Cy3-conjugated anti-rabbit IgG (A102520, Invitrogen), Cy5-conjugated anti-rabbit IgG (A10523, Invitrogen) and Cy5-conjugated anti-rat IgG (A10525, Invitrogen).

### Microscopy and image acquisition

For image acquisition of stained sections, a laser scanning confocal microscope (LSM780 or LSM880, Carl Zeiss) was used equipped with a 40x/1.3 oil DICII PL APO objective. For image acquisition of organoids, we used an inverted spinning disk wide confocal microscope (CSU-W1, Nikon) equipped with a 40x/1.15 CFI APO LWD water objective.

### Single cell RNA-seq analysis

#### Initial mapping, QC and raw counts

3 batches of cells were processed and sequenced using Smart-seq2. FASTQ files were mapped to GRCm38 (mm10) using the STAR aligner (v2.7.7a) (Dobin et al., 2013). Downstream processing was performed using HTSeq (0.13.5) (Anders et al., 2015) to generate the raw gene counts matrices.

Single cell Gene counts and metadata were imported and analyzed in R (v4.3.0) using Seurat (4.3.0.1) (Butler et al., 2018; Satija et al., 2015; Stuart et al., 2019). 3 batches of single cell RNA-Seq data were first processed individually, and then integrated. In batch 2, we filtered out cells from plate 5, since these cells were repeated from Batch1. ENSEMBL gene ids were converted to gene symbols using biomart (v2.56.1), using mouse genome annotation GRCm38, using the following settings: (biomart = ‘genes’, dataset = ‘mmusculus_gene_ensembl’,version = 102) (Durinck et al., 2009). Further analysis was carried out using Seurat.

First, features which were present in >2 cells, and cells having >200 features were selected for analysis. Next, filtering was performed to retain cells with <10%mitochondrial reads, >1400 features (genes) & an RNA count of >100000. The dataset was normalized using Log normalization. Gene expression counts were scaled for all genes. Next, principal component analysis (PCA) was performed using the 2000 most highly variable genes. Clustering was performed using the first 30 principal components (PCs), using a resolution of 0.5. UMAP was created using the first 30 PCs, using a spread of 0.4.

Next, non-mammary-epithelial cell types were filtered. Stromal and salivary cells (cells with gene counts of Epcam<2, Dcpp1>1, respectively), were filtered out. Data normalization, PCA, clustering and UMAP steps above were repeated after cell filtering.

Afterward, integration of all 3 batches was performed using Seurat. During integration, batch 1 was used as a reference, since it had a balanced representation of basal, intermediate, and luminal cell types (determined previously using FACS). After integration, dataset was scaled, and PCA, clustering and UMAP steps were carried out once again on the integrated dataset, using a clustering resolution of 0.5 and UMAP spread of 0.4.

Selection of this cluster resolution was based on the stability of clusters (using tool clustree) (Zappia and Oshlack, 2018), gene expression patterns after clustering (using Seurat’s findMarkers), and FACS cell type labels. 4 clusters were noted. The gene expression markers and FACS labels within clusters were examined. Cluster 1 was composed of basal and intermediate cells (labelled by FACS), hence this cluster was further split (clustering resolution 0.6), giving 2 subclusters: one having a more basal gene expression pattern, and another having a less basal, more intermediate gene expression. Based on expression of known cell type markers and FACS labels, we labelled the 5 resulting clusters as BAS, INT1, INT2, LP and ML. Markers for these clusters were determined using Seurat’s FindAllMarkers tool, using the settings only.pos=TRUE (selecting positive markers), min.pct=0.40 (only test markers which are expressed in at least 40% of cells), and test.use=“roc”. Markers were ordered by Log2 foldchange and the top 7 markers for each cluster were plotted in a heatmap. To estimate expression of basal and luminal gene signatures in each cell, Seurat’s AddModuleScore function was used, using basal and luminal signatures from Kendrick et al. (Kendrick et al., 2008). Additionally, Luminal ER positive and ER negative signatures were combined to give a luminal combined score. Similarly, the cell cycle module scores were calculated using genes from GO and KEGG (GO positive regulation of cell cycle GO0045787, KEGG_CELL_CYCLE.v2023 from Msigdb).

#### Integration / label transfer

Comparison of the Smart-N1ICD with other datasets was performed using label transfer in Seurat. This process allows classification of cells in a query dataset, using another dataset as a reference. Corresponding cell type labels from the reference dataset are transferred to the query dataset, as a ‘predicted ID’. Original clusters (from the query dataset) and corresponding predicted IDs (from the reference dataset) were compared using alluvial plots, by ggplot (https://ggplot2.tidyverse.org), ggalluvial (v0.12.5) (http://corybrunson.github.io/ggalluvial/) (Brunson, 2020).

#### Wuidart dataset

To compare our Smart-N1ICD dataset with the dataset from Wuidart *et al*. (Wuidart et al., 2018), gene counts from Wuidart *et al* were downloaded, and the dataset was re-analyzed and filtered using the same parameters described in Wuidart *et al*., with the following changes, to aid comparison with the Smart-N1ICD dataset: the analysis was performed using Seurat, and Log Normalization was used. Data were scaled, PCA, clustering and UMAP steps were run, using the same settings as for our Smart-N1ICD dataset. Based on gene markers, gene expression clusters were defined as BAS, E14_EMP, and LP and ML.

#### Centonze dataset

The scRNAseq RDS object reported in Centonze *et al*. (Centonze et al., 2020) was downloaded from GEO (GSE148791). To aid comparison with the Smart-N1ICD dataset, the Centonze dataset was re-normalized (using Seurat Log normalization) and re-processed using the same steps described for the Smart-N1ICD dataset. After gene expression clustering, the resulting cell subtypes were re-labelled as BAS, INT_Hyb (intermediate hybrid), LP, ML.

#### SCENIC analysis

To infer transcription factor (TF) activity, SCENIC analysis was performed using pySCENIC (0.12.1) (Aibar et al., 2017; Van de Sande et al., 2020), using default parameters. Smart-seq2 gene expression counts and metadata were imported to create Anndata files (Scanpy, v1.7.2), quality checks were performed and cells were filtered using the same parameters as described for Seurat analysis, to exclude low-quality cells from the analysis. All 3 batches were then concatenated, and then converted to a loom format for analysis with the pySCENIC pipeline. First, genes correlating in expression with TFs were inferred using the GRN step, resulting in TF modules. Next, the CTX step was used to prune genes from these modules, to retain only genes which contain the associated TF motif within cis-regulatory regions. These pruned modules represent regulons of TFs and their associated downstream targets. Lastly, activity of the regulons was estimated as an Area Under Curve (AUC) in the AUCell step of the analysis. From the resulting loom file, a matrix of AUC values was extracted and then imported into an R environment for further analysis. The AUC matrix and the Smrt-NIC Seurat object were both filtered to contain the same set of cells. Next, cluster annotation labels (derived from the Seurat object) were used to perform Regulon specificity score (RSS) analysis on the SCENIC AUC matrix, to infer which regulons show the strongest cluster-specific activity. Activity of selected regulons were also visualized using UMAP plots.

#### Trajectory analysis

Trajectory analysis was performed on the Smart-N1ICD Seurat object using slingshot (v2.8.0) (Street et al., 2018). Slingshot analysis was performed on the PCA structure, specifying the BAS cluster as the origin. Principal curves were plotted and 2 trajectories were observed.

Next, genes whose expression correlated with the trajectory #1 (from BAS to HR^neg^) pseudotime were inferred using tradeseq (v1.14.0) (Van den Berge et al., 2020), which uses a generalized additive model (GAM) to fit the variation of expression of each gene along a pseudotime. The optimal number of knots was estimated for our dataset at 8 knots, which were further used for the analysis. This analysis was performed upon the 4000 most highly variable genes in the dataset. Next, an association test was performed to identify genes correlating with pseudotime, using settings lineages=TRUE and contrastType=”consecutive”. The genes were ordered based on statistical significance of this correlation (Wald statistic), and the top 40 genes significantly associated with trajectory #1 were inferred. To visualize these genes, integrated expression values for these genes were extracted from the Seurat object, ordering cells along trajectory #1 pseudotime. Gene expression values in the resulting matrix were smoothed using a binning/ rolling window process along pseudotime, using the rollapply function from the “zoo” package (1.8-12) (Zeileis and Grothendieck, 2005). Correspondingly, a similar bin smoothing was applied on cluster annotations, selecting the most common cluster annotation within each bin. The resulting matrix was plotted as a heatmap using the pheatmap package (v1.0.12) (https://cran.r-project.org/web/packages/pheatmap/index.html), along with cluster annotations, using settings cutree_rows=3, and clustering_method=”average”, and package viridis (v0.6.3) (Garnier et al., 2024) was used for the heatmap color palette. For selected genes, integrated gene expression was plotted in single cells ordered along the trajectory 2 in pseudotime.

#### Statistical tools

Analysis in R (4.3.0) was performed within RStudio (2023.06.0+421). Analysis in Python (SCENIC) was performed using python (3.10.4) within a conda environment (4.7.12), using Jupyter notebook (6.4.10). ggplot-based plots were created with ggplot2 (v3.4.2).

#### Statistics and reproducibility

Animals were randomized and analyzed in a non-blinded manner. All graphs show mean ± SD. For each experiment, at least n=3 biological replicates were analyzed.

## Data availability

Smart-Seq2 scRNA-sequencing data generated in this study is accessible on the Gene Expression Omnibus GEO repository (GSE268822, available at https://www.ncbi.nlm.nih.gov/geo/query/acc.cgi?acc=GSE268822). The following secure token has been created to allow review of record GSE268822 while it remains in private status: qtozsiowtxunhkj. Analysis codes are available upon request.

## Acknowledgments

We acknowledge Sarah Kinston and Berthold Göttgens at Wellcome Trust-MRC Cambridge Stem Cell Institute, University of Cambridge, UK, for assistance with the Smart-seq2V2 scRNA-sequencing protocol. We also wish to warmly acknowledge the flow cytometry and cell sorting platform at Institute Curie for their technical support, the *in vivo* experimental facility for help in the maintenance and care of our mouse colony, and the Cell and Tissue Imaging Platform-PICT-IBiSA at Institut Curie (member of the French National Research Infrastructure France-Bioimaging, ANR-10-INBS-04) for their expertise and assistance. We are very grateful to all members of the Fre laboratory for support, critical reading of the manuscript and constructive discussions. This work was funded by Paris Sciences et Lettres (PSL* Research University) (grant # C19-64-2019-228), the French National Research Agency (ANR) grant numbers ANR-21-CE13-0047 and ANR-22-CE13-0009, the Medical Research Foundation FRM “FRM Equipes” EQU201903007821, the FSER (Fondation Schlumberger pour l’éducation et la recherche) FSER20200211117, the Association for Research against Cancer (ARC) label ARCPGA2021120004232_4874, the Worldwide Cancer Research Foundation # 24-0216 and by Labex DEEP ANR-Number 11-LBX-0044 to SF. C.M. was funded by a post-doctoral fellowship from ARC (ARCPDF12021020003033).

The funders had no role in study design, data collection and analysis, decision to publish, or preparation of the manuscript.

